# Comparative genomics groups phages of Negativicutes and classical Firmicutes despite different Gram-staining properties

**DOI:** 10.1101/606087

**Authors:** Chris M Rands, Harald Brüssow, Evgeny M Zdobnov

## Abstract

Negativicutes are Gram-Negative bacteria characterized by two cell membranes, but they are phylogenetically a side-branch of Gram-Positive Firmicutes that contain only a single membrane. We asked whether viruses (phages) infecting Negativicutes were horizontally acquired from Gram-Negative Proteobacteria, given the shared outer cell structure of their bacterial hosts, or if Negativicute phages co-evolved vertically with their hosts and thus resemble Gram-Positive Firmicute prophages. We predicted and characterised 485 prophages from Gram-Negative Firmicute genomes plus 2,977 prophages from other bacterial clades, and we used virome sequence data from 183 human stool samples to provide independent evidence for our predictions. The majority of the identified Negativicute prophages were lambdoids closer related to prophages from other Firmicutes than Proteobacteria by sequence relationship and genome organization (position of the lysis module). Only a single Mu-like candidate prophage and no clear P2-like prophages were identified in Negativicutes, both commonly found in Proteobacteria. Therefore, it is unlikely that Negativicute phages were acquired from Proteobacteria. Sequence-related prophages were identified in two distinct Negativicute orders (Veillonellales and Acidaminococcales) possibly suggesting horizontal cross-order phage infection between human gut commensals. Phages infecting these Negativicute orders occasionally harboured putative antibiotic resistance genes.

## Introduction

The number of bacterial and archaeal species, or so-called operational taxonomic units, has been estimated at between 0.7 and 4.3 million using 16S rRNA sequences (Louca et al., 2019). Despite frequent horizontal gene transfer, phylogenetic trees can still be developed for bacteria, allowing sequence-based taxonomic classification. Bacterial virus or bacteriophage (phage) species might outnumber bacterial species since most investigated phages are host-specific and most bacterial species are infected by several distinct phages. The high percentage of unknown genes identified in metavirome analyses supports the notion that phages are very diverse (Yutin et al., 2018). Although rampant horizontal gene transfer between phages was predicted by the modular theory of phage evolution (Botstein, 1980), comparative genomics has revealed aspects of vertical evolution in bacteriophages and ancient lineages. Phages of a given type share substantial sequence homology when infecting the same bacterial species or genera, but for increasingly distant bacterial hosts, the ability to identify homology gradually decreases until the phage sequences can no longer be aligned. However, conservation of aspects of phage genome organization (i.e. gene order) was observed between phages infecting distantly related hosts, for example across lambdoid phages from Proteobacteria, Firmicutes and viruses infecting Archaea (Lucchini et al., 1998; Pfister et al., 1998; Desiere et al., 1999; Lucchini et al., 1999; Desiere et al., 2000; Brüssow and Desiere, 2001; Casjens and Hendrix, 2015; Kang et al., 2017; Mahmoudabadi and Phillips, 2018). Structural biologists even detected relatedness between the protein folds of capsid components from phages and animal viruses (Bamford et al., 2005). These observations indicate that phage lineages have a very ancient shared ancestry and diversified in parallel with their host bacteria through co-evolution.

In general, bacterial lineages accumulate small phenotypic (i.e. trait) changes, making close bacteria-phage co-evolution feasible. However, occasionally large bacterial phenotypic changes occur (Cavalier-Smith, 2006), potentially presenting a challenge for their phages to adapt. One such example comes from the Negativicutes. Bacterial genome sequence analysis places Negativicutes (prototype *Veillonella parvula*) within the Firmicutes (Marchandin et al., 2010; Campbell et al., 2015), a clade of almost exclusively Gram-Positive bacteria (prototype *Bacillus subtilis*) with a single membrane and a thick peptidoglycan layer. However, Negativicutes are clearly Gram-Negative bacteria since they possess an inner cell membrane, a thin peptidoglycan layer and an outer membrane with lipopolysaccharides. In this respect Negativicutes resemble Gram-Negative Proteobacteria (prototype *Escherichia coli*). Therefore, during evolutionary history Firmicute phages had to adapt to a different membrane boundary, assuming vertical phage-bacteria co-evolution. An alternative is that Negativicute phages were acquired horizontally from Proteobacteria, plausible given the shared outer cell structure of their bacterial hosts, and possible environmentally since both bacterial groups share an ecological niche in human microbiomes, particularly in the human gut (Rands et al., 2018).

We used the unusual relationship between Negativicutes and classical Gram-Positive Firmicutes to investigate phage-bacteria co-evolution. Apart from the Negativicutes, the only other Gram-Negative Firmicute group are the Halanaerobiales, a very poorly sampled bacterial order whose members inhabit saline environments like lakes. Phylogenetically Negativicutes are closest to Gram-Positive Clostridiales, while Halanaerobiales are closest to Gram-Positive Natranaerobiales, so the two Gram-Negative clades form independent branches of Firmicutes (Antunes et al., 2016). If phages co-evolved in parallel with their bacterial hosts, we would expect Negativicute and Halanaerobiales phages to be closer to phages from Gram-Positive Firmicutes that those of Proteobacteria. Conversely, we might find Negativicute phages are grouped with phages from Proteobacteria, “attracted” by the similar cell surface structure, since lipopolysaccharides and outer membrane proteins are the primary point of contact for phages in the infection process. These hypotheses have not been tested since sequence data on phages from Gram-Negative Firmicutes is currently limited to a few incomplete Negativicute phage sequences reconstituted from human and mouse virome projects (Pride et al., 2012; Kim and Bae, 2016), some predicted but uncharacterised prophages (Roux et al., 2015), and two possibly defective lambdoid prophages (Rands et al., 2018).

Here we predicted and characterised 485 prophages from Gram-Negative Firmicute genomes and compared them to 2,977 prophages that we predicted in other bacterial lineages. We found that both the genome organization and sequence-relatedness of lambdoid Negativicute and Halanaerobiales prophages identified them as closer relatives of lambdoid phages from Gram-Positive Firmicutes than prophages from Proteobacteria. Phages frequently found in Proteobacteria, like Mu-like and P2-like prophages, were only very rarely or not at all detected in Negativicutes. These observations are not expected under a hypothesis of substantial phage gene flow between Proteobacteria and Gram-Negative Firmicutes.

## Results and Discussion

### Bacterial genome screening identifies prophages from Gram-Negative Firmicutes

As a first step towards an evolutionary analysis of Gram-Negative Firmicute phages, we screened all available Negativicute and Halanaerobiales bacterial genomes deposited in the NCBI Genbank database for sequences of prophages. Using PHASTER (Arndt et al., 2016) with additional post-processing and annotation steps (see Methods and materials) we predicted and annotated 485 prophages across 350 Gram-Negative Firmicute bacterial genome assemblies. For comparison, we also predicted prophages in 200 complete bacterial genomes from each of *Escherichia* and *Bacillus*, and 136 complete genomes (the total available) from *Clostridium*.

Among Gram-Negative Firmicute phages, 41% of the predicted prophages were from the Negativicute order Selenomonadales (genera *Selenomonas* and *Sporomusa*), 39% were from the Negativicute order Veillonellales (genera *Veillonella* and *Megasphaera*), 8% from the Negativicute order Acidaminococcales (genera *Acidaminococcus* and *Phascolarctobacterium*), and 7% from various other Negativicute taxa. Five per cent of prophage sequences came from the order Halanaerobiales (genus *Halanaerobium*) (**Figure S1**, **Table S1**). For genera represented with more than 8 sequenced bacterial genomes, the highest viral frequency was 4.1 prophages per bacterial genome, found in *Sporomusa*, while the lowest frequency of 0.3 prophages per bacterial genome was detected in *Halanaerobium*. Overall, a mean of 1.5 prophages per sequenced Negativicute bacterial genome was detected. This figure was only slightly higher, namely 1.9, when calculated for completely sequenced and assembled Negativicute genomes. A typical bacterial genome contains around 3 prophages based on *in silico* prediction (Kang et al., 2017), similar to our estimate of 3-4 prophages per genome for *Clostridium* and *Bacillus* (**Table S2**), while higher values are normal for Gram-Negative *Enterobacteriaceae* (Bobay et al., 2014), indeed, we estimate 8.8 prophages per *Escherichia* genome. However, the low prophage content in Negativicutes and particularly in Halanaerobiales should be interpreted with caution: prophages might be missed since sequenced phages from Gram-Negative Firmicutes are very few and protein annotations are lacking, making prophage detection more difficult computationally. Prophage proteins lacking any sequence homology to previously known phage proteins will likely escape detection.

### Virome screening for Negativicute phages confirms prophage predictions

To circumvent this difficulty of a deficit of annotated phage proteins, we used viral-like particle (VLP) (“virome”) data to confirm that we did not miss obvious prophage predictions. We screened 183 samples of virome sequences from human stool for matches to four completely sequenced strains of Negativicute that are residents of the human gut. The strains were: *Acidaminococcus intestini* RYC-MR95, *Veillonella parvula* UTDB1-3, *Dialister Marseille*-P5638, and *Negativicoccus massiliensis* Marseille-P2082, all predicted to contain at least one prophage (**Table S3**). We processed and aligned the virome reads to these bacterial genome sequences (see Materials and Methods). If the VLP sequences contain no temperate phages infecting these bacteria, we will find only few virome reads aligning with the bacterial genome, likely representing bacterial DNA contaminating the viral fraction or bacterial DNA packaged into the phage capsid by generalized transduction. If the virome read alignments overlap the predicted prophage sequences, the prediction is independently confirmed. It is, however, also possible that a predicted prophage lacks a virome counterpart, which suggests a prophage remnant that is not any longer active in producing an extracellular phage or a phage simply not present in the examined sample. The virome sequences confirmed the previously described prophages 1 and 2 in *A. intestini* (Rands et al., 2018) and prophage 4 we predicted here (**Figure S2A**). The identified further sharp peak in read depth at around 700kb, likely represents an erroneously packaged element like a small integrative plasmid or conjugative transposon (**Table S4**), similar to a previously reported element (Warburton et al., 2009).

For *V. parvula* the two predicted prophages were confirmed (**Figure S2B**), and two further peaks represent bacterial DNA (**Table S4**). For *Dialister*, few reads match the predicted prophage and another small peak covers one kb of unannotated DNA (**Table S4**, **Figure S3A**). For *Negativicoccus*, five genome regions showed weak matches with virome sequences (**Figure S3B**), only one corresponds to a predicted prophage. Two larger peaks correspond to an Illumina PCR Primer and bacterial DNA (**Table S4**). Overall, none of the read alignment peaks not overlapping a prophage prediction appear to correspond to real phages. Therefore, while homology-based computational prediction of prophages from bacterial genome sequences may underestimate the true prophage content by missing highly divergent phages, the read alignment of virome data to bacterial chromosomes does not suggest that we missed clear prophages in these four test cases.

### A common genome organization among prophages from Negativicutes and Gram-Positive Firmicutes

Next, we analysed the genome organization of the predicted Gram-Negativicute Firmicute prophages. We annotated the prophage genes and color-coded them according to their putative function and constructed prophage genome maps. Alignment of the genome maps showed a common pattern of genome organization for the best annotated prophage predictions from Negativicutes. Over the DNA packaging, head and head to tail genes, the gene organization was striking conserved and shared a similar gene order with lambdoid prophages from both Firmicutes and Proteobacteria (Figure 1). Halanaerobiales prophages also showed the typical genome map of lambdoid prophages, although few complete prophages were predicted, due to the paucity of available bacterial genomes. We conducted a gene flanking analysis with the entire predicted prophage set by asking what annotated genes are found adjacent to the query gene. We found that integrase genes were most frequently flanked by phage lysogeny genes (e.g. phage repressors) on one side and bacterial genes on the other side. DNA replication genes (primases and helicases) were flanked by lysogeny and DNA packaging genes (terminases), respectively. The phage terminase gene had frequently another terminase gene on one side and a portal protein gene on the other side. The prohead protease or scaffold genes were bracketed by portal and head genes. Head-to-tail genes were found next to head genes on one side and tail genes on the other side. Tail sheath genes were flanked by head-to-tail and tail genes. Tail, tail tube, and tail tape measure genes showed mostly other tail genes next to them. Notably, tail fibre genes were next to other tail genes on one side and holin or lysin or lysogeny genes at the other side (Figure 1).

**Figure 1:**
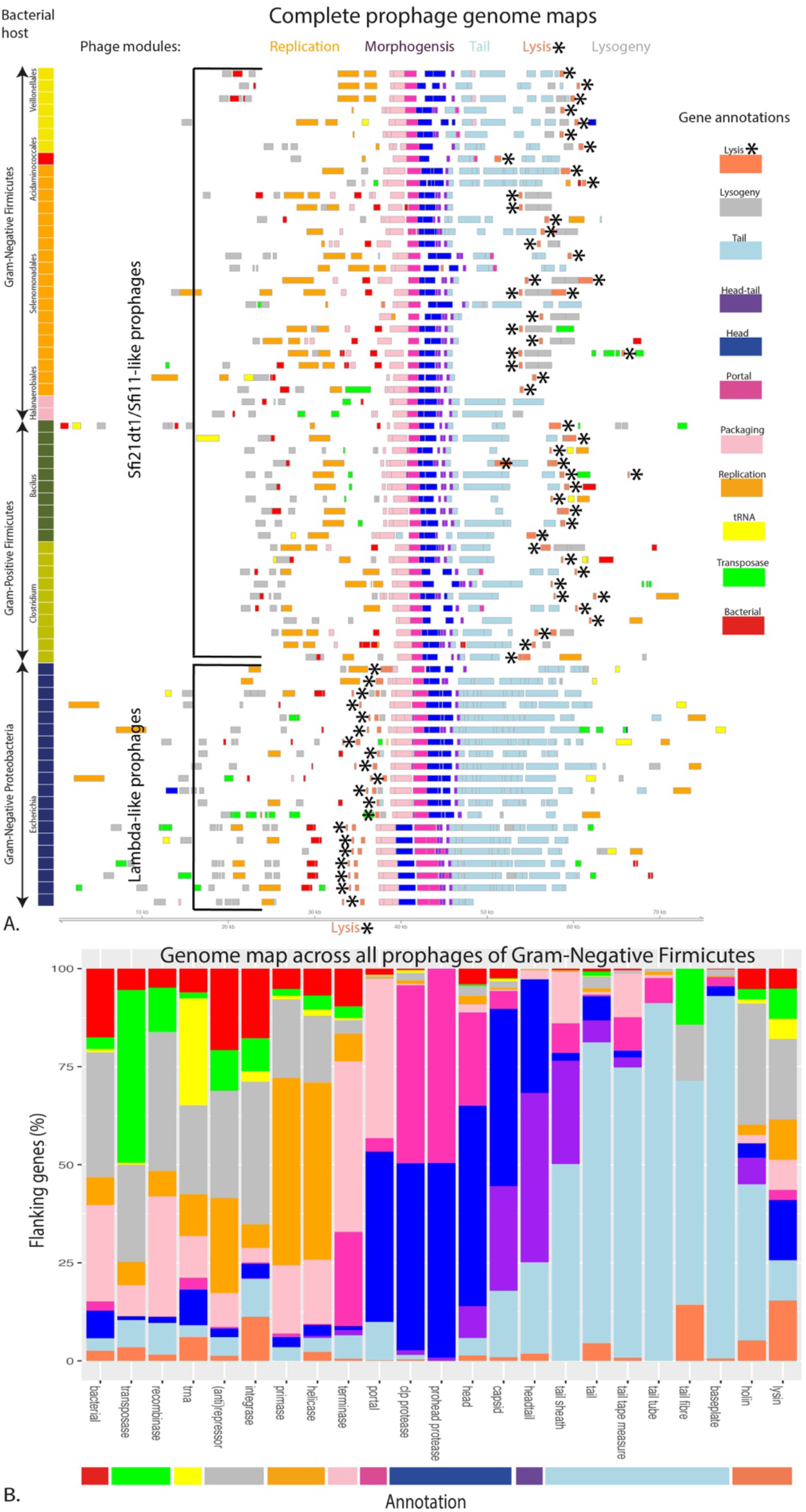
Genome organization of prophages. **A.** Genome maps of the most complete and best annotated lambdoid prophages predicted from 350 Gram-Negative Firmicute genomes (**Table S1**) and examples predicted from the genomes of prophages from the genera *Clostridium, Bacillus* (Gram-Positive Firmicutes) and *Escherichia* (Gram-Negative Proteobacteria). Each line depicts an individual prophage and each rectangle represents a predicted gene with length and prophage genome position to scale. The genes are color-coded according to the functional annotation of different phage modules as specified in the top right colour key. The left vertical colour scale identifies the taxonomic attribution of the host bacteria. All these prophages show the typical genome organization of lambdoid phages as most evident over the morphogenesis gene modules with a typical gene order (see text for details; exception: Enterobacteria phage 2851 / Stx2-converting phages of Escherichia, shown at the bottom, that have swapped the capsid and portal gene positions). A characteristic difference is found between lambdoid prophages from Firmicutes (both Gram-positive and Gram-negative) and Proteobacteria in the position of the lysis module (highlighted by *). **B.** Gene flanking analysis of prophage genes. For each phage protein that was annotated among the 485 predicted prophages from Negativicutes we took the nearest annotated neighbour on each side of that protein and tallied up their frequencies.

This order across the morphogenesis genes mirrors that of typical lambda-like prophages from Firmicutes (Lucchini et al., 1999). However, the phage lysis module (holin, lysin genes) has a different genome position in lambdoid prophages from Proteobacteria (Figure 1A). In Firmicute lambdoid phages the lysis cassette follows directly the tail genes and precedes the lysogeny module as observed for *Streptococcus* Sfi21dt1/Sfi11-like phages (Canchaya et al., 2003b). However, in lambdoid prophages from Proteobacteria the lysis module is located after the phage DNA replication module and precedes the DNA packaging (terminase) genes like the classical Lambda phage. This distinct positioning of the lysis module in phages from Firmicutes and Proteobacteria is apparently a general rule. There are no exceptions looking at the best annotated prophage genome maps (Figure 1A), and the same trend is observed when considering all prophages (**Figure S4)**. Rare exceptions to the rule may be due to mis-annotations, prophage remnants, recombination events or non-lambdoid prophages and none of the few candidate Negativicute phage exceptions represented convincing examples of intact lambdoid phages upon further inspection of the genome maps and protein annotations. We are aware of only a single exception from isolated sequenced phages, *Pseudomonas* phage D3 infecting a Proteobacterium displays the Firmicute prophage-specific lysis gene constellation (Kropinski, 2000). It is unlikely that this different lysis cassette positioning reflects distinct transcriptional regulation of the phage lysis program necessitated by the presence of two vs. one host membranes that must be crossed during progeny phage release, since the prophages from Negativicutes egress from bacteria enclosed by two membranes while maintaining the lysis module position of Firmicutes. The positioning of the lysis cassette might just represent a conserved trait inherited from ancestor phages, but it remains a useful evolutionary marker for phylogenetic studies.

Negativicute phages are thus linked by their prophage genome maps with classical Gram-Positive Firmicute phages, inconsistent with widespread acquisition of Negativicute phages from Proteobacteria and instead supporting the vertical co-evolution model. In this evolutionary scheme one would also expect few Mu-, P2- or Inovirus-like prophages in Negativicutes, which are commonly found in Proteobacteria. We found only a single Negativicute prophage that displayed similarity to a Mu-like prophage genome map (~50kb size with a characteristic gene order: two transposition genes, transcriptional activator, lysis, head and tail genes comprising baseplate genes, followed by a reverse transcriptase genes) (**Figure S5**). However, this observation is not without precedence since similar Mu-like prophages were occasionally found in Gram-Positive Firmicutes (Toussaint, 2013). We did not detect clearly P2-like prophages in Negativicutes. However, this negative result must be interpreted with caution since we will miss prophages that lack homology to known phage annotated proteins and are absent from virome sequences. Our methods are not designed to detect Inoviruses, but a recent study did not find evidence of these in Negativicutes, although they sampled relatively few Negativicute genomes, and unexpectedly they did report possible Inovirus-like prophages in Gram-Positive Firmicutes (Roux et al., 2019).

### Sequence relatedness of the Negativicute phages identifies evolutionary relationships among prophages

In the next step we identified homologous taxonomic groups of phages based on sequence information (see Materials and Methods). Across the full prophage genome sequences, the phages of Gram-Negative Firmicutes, Gram-Positive Firmicutes, and Proteobacteria always formed separate groups. This is indicated in Figure 2A,B where individual phages are depicted as dots coloured by their bacterial host with each grid reference showing a different cluster of phages. The clusters with large number of phages appear as blocks of homogenous colour so the individual dots are not visible. At the 80% nucleotide sequence identity connections were, with one exception, limited to prophages infecting the same taxonomic order. The exception was high nucleotide sequence sharing between a prophage pair retrieved from Veillonellales and Acidaminococcales (Figure 2B, grid reference A1), two different orders of Negativicutes. When the threshold was decreased to 40% nucleotide sequence identity over the prophage genome, this number increased to four pairs of related Veillonellales and Acidaminococcales phages (Figure 2A, grid reference G3). In fact, when Veillonallales and Acidaminococcales prophages were classified according to sequence relationship within this cluster, no correlation with the bacterial host order was observed suggesting frequent phage exchanges between these two Negativicute groups (Figure 3). The high nucleotide sequence identity between prophages from two different orders of bacteria is unusual. This close relationship between Veillonellales and Acidaminococcales prophages might not necessarily reflect close evolutionary relationships of the bacterial hosts but could instead be the result from cross-order phage infection. With laboratory-adapted phages, infections across genus borders are rare and sequence homology analyses generally support this (Gao et al., 2017), but microbial ecologists have recently provided data suggesting that phage infections across larger taxonomic borders occur in the natural environment at much higher frequency than anticipated from laboratory experiments (Kauffman et al., 2018).

**Figure 2:**
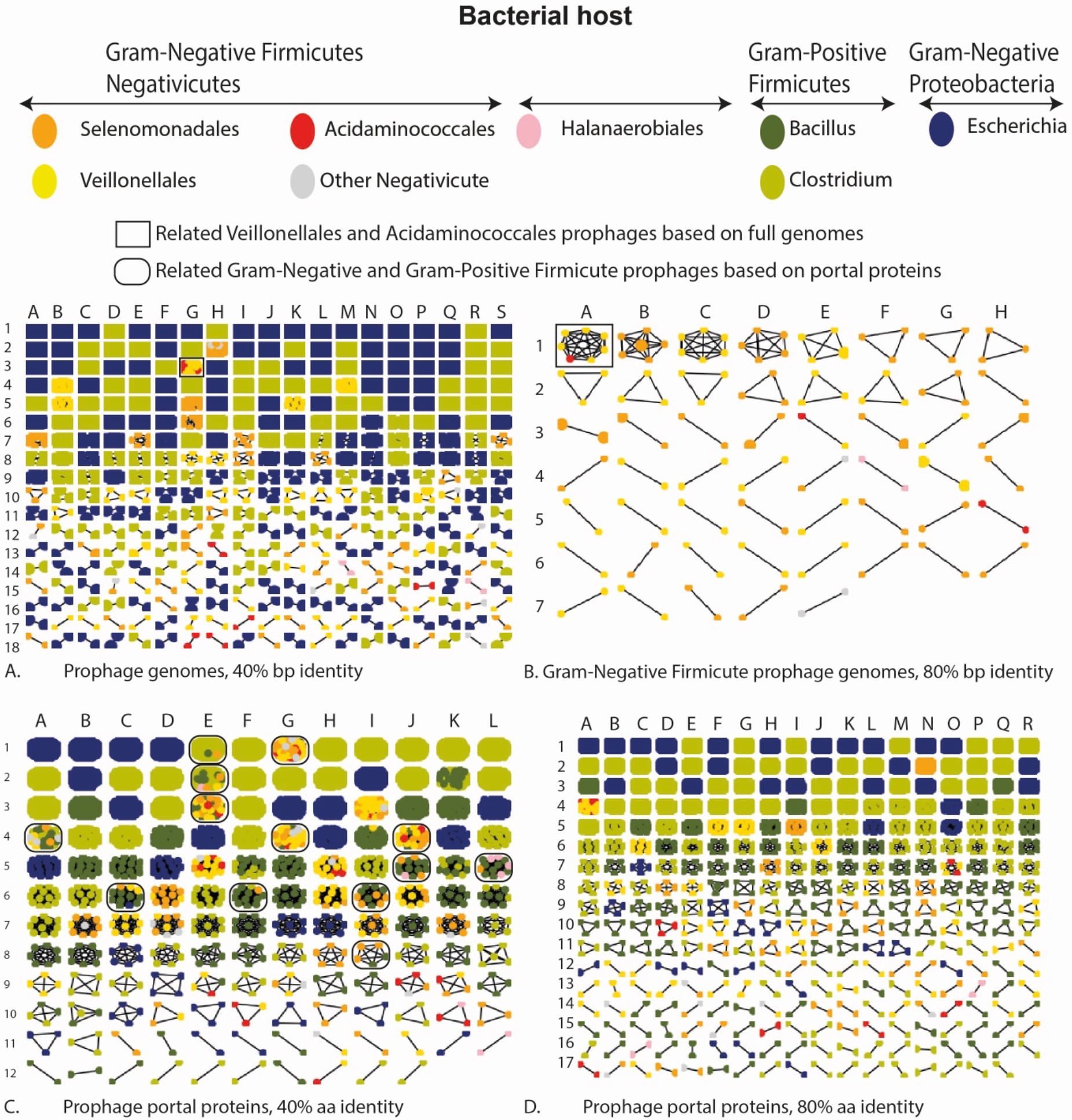
Sequence relationships among prophages using all 485 predicted phages of Gram-Negative Firmicutes and 2,977 other prophages predicted from other Firmicutes and Proteobacteria (**Tables S1, S2**). Each dot represents an entire prophage genome (panels **A**, **B**) or only the prophage portal protein (panels **C**, **D**) coloured with respect to the bacterial host. Each grid reference represents a group of phages that cluster together at the indicated nucleotide (bp) (panels A, B) or protein (aa) (panels C, D) sequence identity threshold. Phages showing no homology to other sequences are not shown (i.e. single dots are omitted). When the dots are highly dense a solid block of colour shows a larger group of sequence-related prophages infecting the same bacterial host.

**Figure 3:**
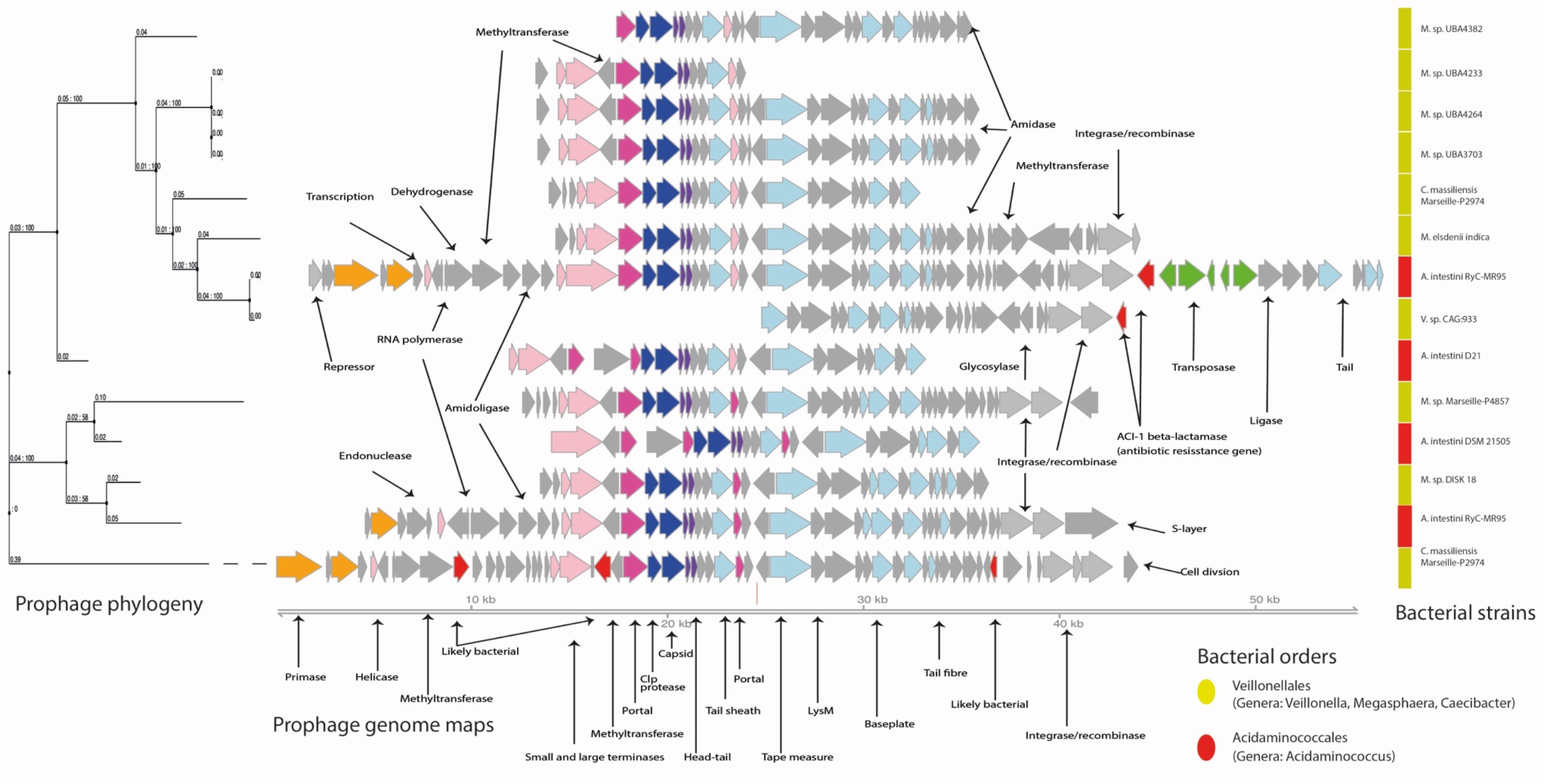
A cluster of Negativicute prophages sharing sequence homology (corresponding to Figure 2A grid reference G4). Shown are the genome maps (center), annotated genes are labelled, phage modular attribution is coloured, see key in Figure 1, red: the previous described *aci1* gene (Rands et al., 2018) and other likely bacterial genes. Displayed according to the Maximum-Likelihood bootstrapped phylogeny based on the full-length prophage sequences (left tree). The right vertical column indicates the bacterial hosts, coloured by Negativicute order.

Neither Veillonellales nor Acidaminococcales prophages shared nucleotide homology with Selenomonadales prophages (Figure 2A,B), the other large order of Negativicutes. The position of Selenomonadales is not certain in bacterial phylogenetic analysis, some authors put them closer to Veillonellales (Antunes et al., 2016), others place them closer to Acidaminococcales (Campbell et al., 2015), while others left the phylogenetic relationship of the Negativicute orders unresolved (Vesth et al., 2013; Yutin and Galperin, 2013). Prophage sequence relationships suggest a closer relationship between Veillonellales and Acidaminococcales to the exclusion of Selenomonadales. However, it is unclear whether prophage sequences can be used as markers for resolving phylogenetic relationships of their host bacteria.

When restricting the analysis to a single relatively conserved protein, the portal protein, we observe that prophages of Negativicutes shared 40% amino acid sequence identity with Halanaerobiales and Firmicute prophages, but not with portal proteins from prophages of Proteobacteria (Figure 2C, e.g. grid references A4, E1, E2, E3, F6, G1, G4, I6, I8, J4, J5 and L5) with only a single exception (grid reference C6, which contains just one *E. coli* phage portal protein with portal proteins of Gram-Positive and Gram-Negative Firmicute prophages). This pattern of sequence conservation concurs with the genome map analysis in identifying the closer evolutionary relationship between Negativicute and classical Firmicute prophages to the exclusion of Proteobacteria prophages. Negativicute prophages are thus clearly not derived from Proteobacteria prophages, so prophage analysis thus suggests that the phage and bacterial phylogenies are concordant, despite the different Gram-staining properties of the bacterial host, and consistent with a model of vertical phage-bacteria co-evolution.

### Negativicute prophages occasionally harbour antibiotic resistance genes

It is disturbing that a prophage from *Acidaminococcus* that is related to other prophages in different bacterial orders (Figure 3) was shown to be associated, in a transposon context, with an antibiotic resistance namely the aci1 gene encoding a class A beta-lactamase (Rands et al., 2018), which confers resistance to penicillins and extended-spectrum cephalosporins (Galán et al., 2000). Here we also found that three different *Veillonella* species carried genes annotated as metallo beta-lactamase (MBL) genes in a prophage map context. The genome maps are conserved and across the full prophage sequences the mean pairwise sequence identity is 49%. However, the prophage context is not clear since there is no DNA replication module and there are several putative plasmid annotated proteins (**Figure S6**). Additionally, while MBLs are known to provide resistance to carbapenems, they have also diverse other functions as hydrolytic enzymes. Until the antibiotic resistance function is biologically demonstrated for these *Veillonella* prophage genes, its possible resistance function should be interpreted with caution. Notably, one *Selenomonas* prophage element contained a better characterised antibiotic resistance gene, tetracycline resistance gene *tet(32)*, related to one found in a *Clostridium* species (Melville 2001). This ARG is in a clear prophage remnant that has suffered substantial rearrangement, but the flanking integrases could still confer mobility to the *tet(32)* gene (**Figure S6**). The *tet*(32) gene was found in a small integrative plasmid or conjugative transposon context from the virome data (**Table S4**).

Our results are of public health interest given the current antibiotic resistance crisis since Negativicutes are common human gut commensals, potentially providing a reservoir of ARGs that might spread to pathogens. Transposons, plasmids and phages are all mobile elements that can facilitate the lateral transfer of ARGs among bacteria, but it remains controversial how frequently phages encode ARGs (Enault et al., 2017). Previous analyses identified a transposon-associated ARG, *aci1*, in an *Acidaminococcus* prophage (Rands et al., 2018). Here we revealed another putative ARG, MBL, in a possible context of a *Veillonella* prophage and another probable ARG, tet(32), in a rearranged *Selenomonas* prophage remnant. So, unambiguous cases of Negativicute prophages encoding and likely mobilising ARGs are apparently uncommon.

### Comparison of prophages and extracellular Negativicute phages

In two faecal samples from human subjects with inflammatory bowel diseases the phage sequence abundance was high enough to generate an assembly of two free extracellular Negativicute phages from the virome sequences. One prophage - extracellular phage pair from *Veillonella parvula*, whose head and lysis gene constellation attributed them to the Sfi11like (Brussowvirus) lineage of lambdoid Firmicute phages, shared highly related DNA replication and transcriptional regulation modules (Figure 4A). Sequence relatedness dropped substantially over the lysis and lysogeny module and the tail fiber region and returned to moderately elevated levels over the remainder of the virion structural genes. This type of modular phage genome organization mirrors closely that observed in streptococcal phages from Firmicutes (Canchaya et al., 2003b). Another prophage - extracellular phage pair from *V. parvula*, whose characteristic portal-Clp protease-major head gene constellation attributes them to the Sfi21dt1virus (Moineauvirus) lineage of lambdoid Firmicute phages, shared high sequence identity over the virion structural genes, but the non-structural genes are absent, implying a prophage remnant unable to induce (Figure 4B).

**Figure 4:**
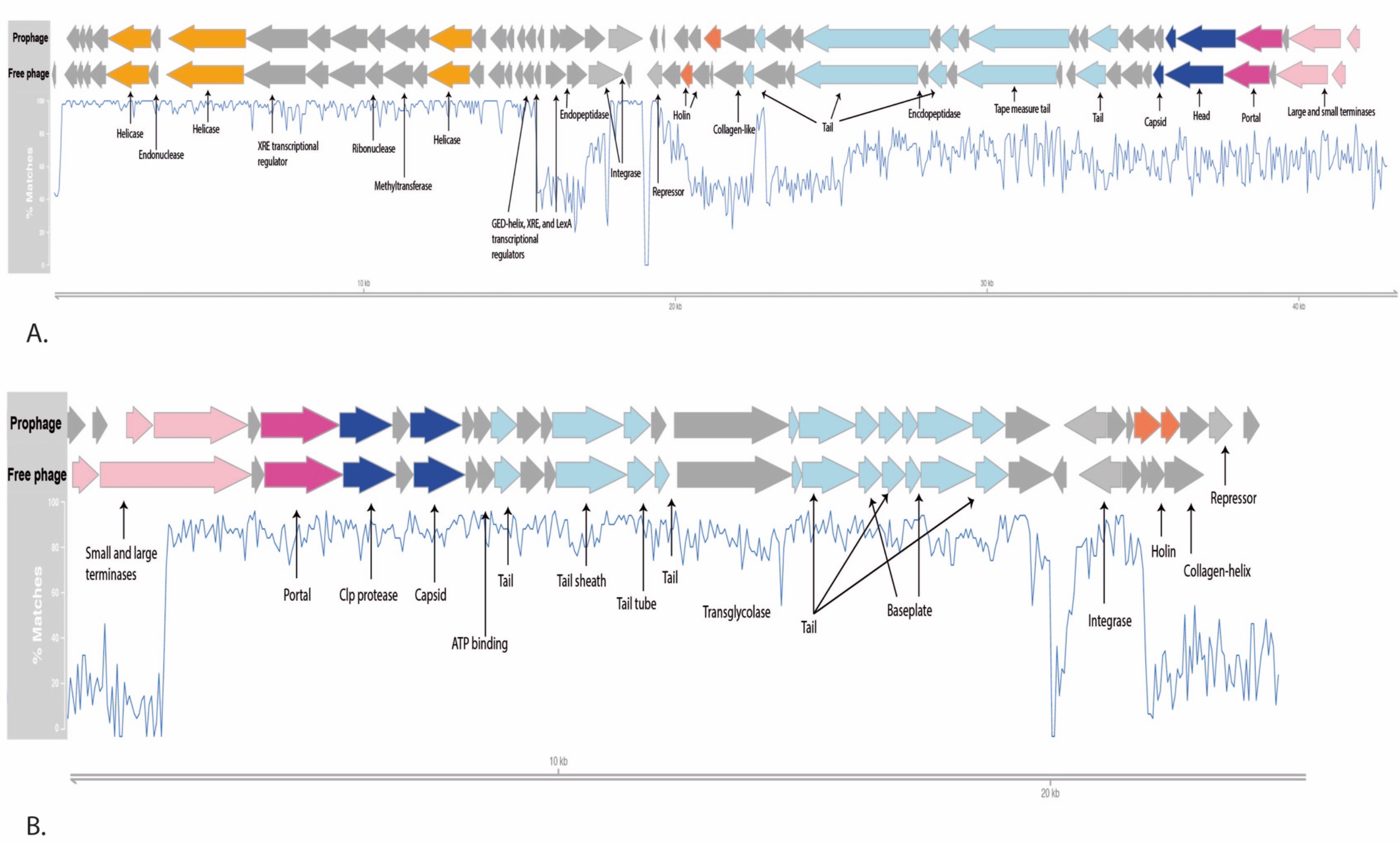
Comparison of the genome maps of two *Veillonella parvula* UTDB1-3 prophages (panel **A**: prophage 1, Sfi11-like lineage, panel **B**: prophage 2, Sfi21-like lineage, see **Figure S2B**) and their related assembled extracellular virus counterparts (free phage) from the fecal virome of two inflammatory bowel disease patients. Phage genes are coloured according their modular attribution (legend Figure 1) and key genes are annotated. The lower blue continuous line gives the percentage matches based on global alignment of the phage genome sequences.

### Chromosomal location of Negativicute prophages in bacterial genomes is not random

Prophage integration has a profound impact on bacterial chromosome structure and bacterial evolution (Canchaya et al., 2003b; Brüssow et al., 2004; Bobay et al., 2013). Complete Negativicute sequences assembled into a circular bacterial genome allow the localization of the prophage insertion sites with respect to the bacterial origin and terminus of replication. These bacterial genomes replicate bidirectionally and the regions near the origin generally have highly transcribed genes due to gene dosage effects. Since prophages are transcriptionally mostly silent and should not interfere with the expression of bacterial household genes, it was postulated that prophages should preferentially be inserted away from the origin of replication (Canchaya et al., 2004; Bobay et al., 2013; Oliveira et al., 2017). Indeed, dividing up Negativicute genomes into twenty equally sized windows, with each window representing 5% of the chromosome length, we found that Negativicute prophages were not distributed randomly across bacterial genomes (p<0.01, χ^2^ test), with a trend towards insertion towards the terminus of replication (Figure 5).

**Figure 5:**
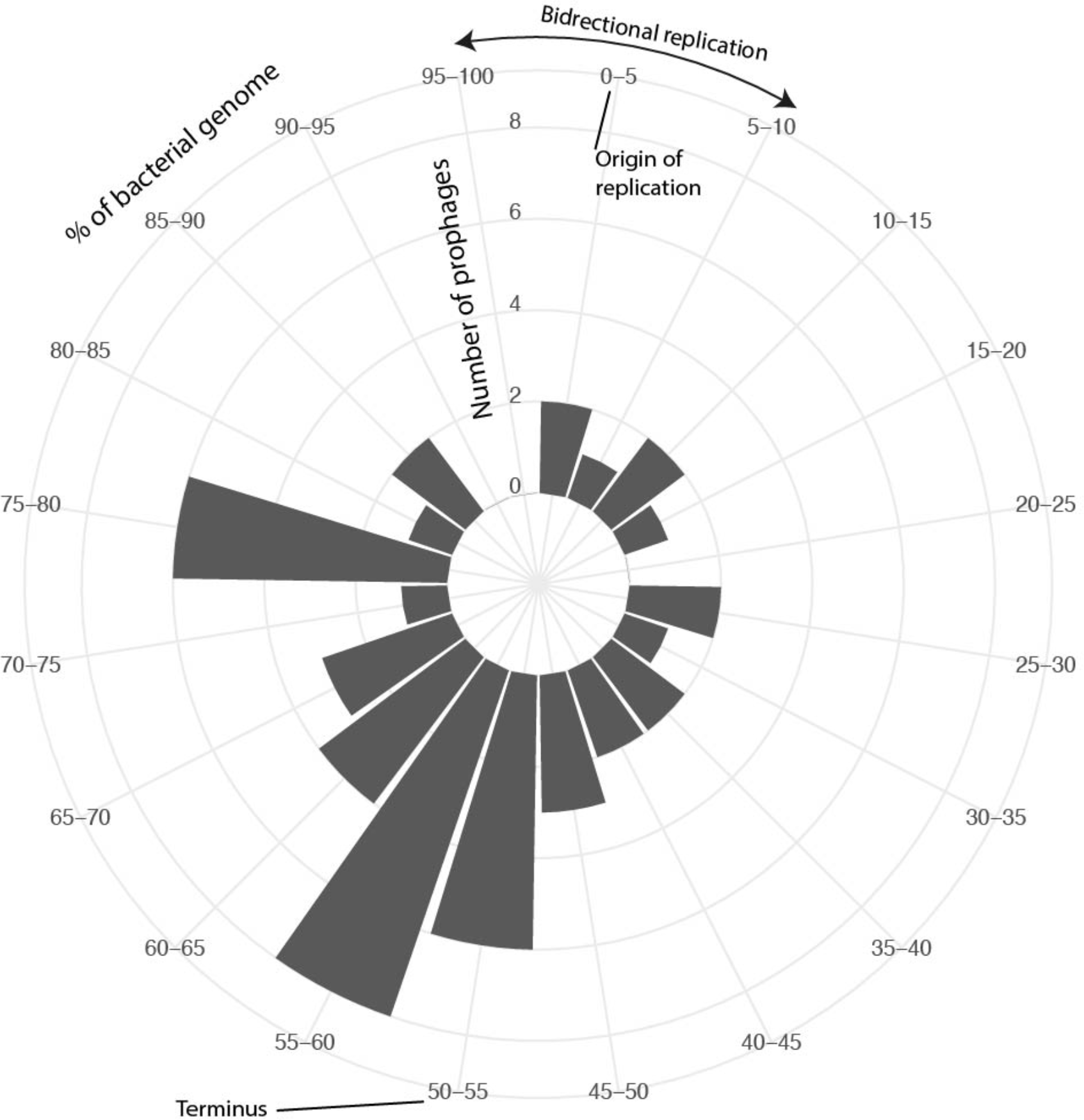
Distribution of prophages insertion sites as bar charts (concentric circles indicate the number of observations) projected on the complete bacterial genomes from Gram-Negative Firmicutes with respect to the origin and terminus of bacterial replication. Bacterial genomes were normalized by size and divided up into windows representing 5% of each genome size on the perimeter of the outermost circle.

### Trends in Negativicute prophage decay

The mean prophage genome size of 24kb across Negativicute and Halanaerobiales prophages is too small for complete lambdoid Firmicute prophages. In fact, only about 10% of the prophages showed a genome length larger than 40kb, the standard genome size of lambdoid prophages. Even among complete finished Negativicute genome assemblies the mean prophage genome size is only 27kb, implying that fragmented prophage predictions are not the main reason for this observation. An alternative explanation is that prophages degrade by gene deletion to become prophage remnants (Bobay et al., 2014). It has been argued that the decay of prophage genomes may not be random. Genes encoding lysogenic conversion functions, transcriptional regulators or recombinases mostly located in the lysogeny module, could be of selective value to the bacterial fitness resulting in their preferential maintenance in prophage remnants (Canchaya et al., 2002; Canchaya et al., 2003a; Bobay et al., 2014). However, for Negativicutes, we did not find a clear correlation between prophage genome size and the functional modules preserved, so these prophages appear to decay with comparable probability across the prophage genomes as they become remnants, although the trend is noisy (**Figure S7**). One might speculate that for the currently sequenced Negativicutes, fitness factors are not frequently mobilized by the phage integrase. Indeed, although we found a few phage encoded ARGs discussed above, there were no clear examples of Negativicute prophages encoding bacterial virulence factors based a scan of the VirulenceFinder database (see Materials and Methods), which contrasts with known examples in pathogenic Firmicutes and Proteobacteria (Brüssow et al., 2004).

### Conclusions

Our study demonstrates that bacterial genome sequences combined with virome sequence data are valuable resources for comparative phage genomics, provided that bacterial genomes and viromes are derived from the same ecological context. We provide insights into the co-evolution of Gram-Negativicute phages and their bacterial hosts. We found evidence of possible cross-order infection between Veillonellales and Acidaminococcales phages, both common human gut commensals, which might also indicate that these two bacterial orders are closer relatives to each other than to Selenomonadales. Of public health relevance, predicted prophages infecting these bacterial orders occasionally harboured putative antibiotic resistance genes. More fundamentally, we showed that prophages from Gram-Negative Firmicutes are closer related to prophages from Gram-Positive Firmicutes than to those of Proteobacteria, based on both the position of the lysis module in the genomes of lambdoid prophages and sequence relatedness among phage portal proteins. There is also an apparent rarity of prophage types in Negativicutes that are common in Proteobacteria. Therefore, Negativicute phages appear to have co-evolved vertically with their Firmicute hosts and not made the horizontal leap from Proteobacteria despite the shared outer membrane structure of their bacterial hosts (Figure 6).

**Figure 6:**
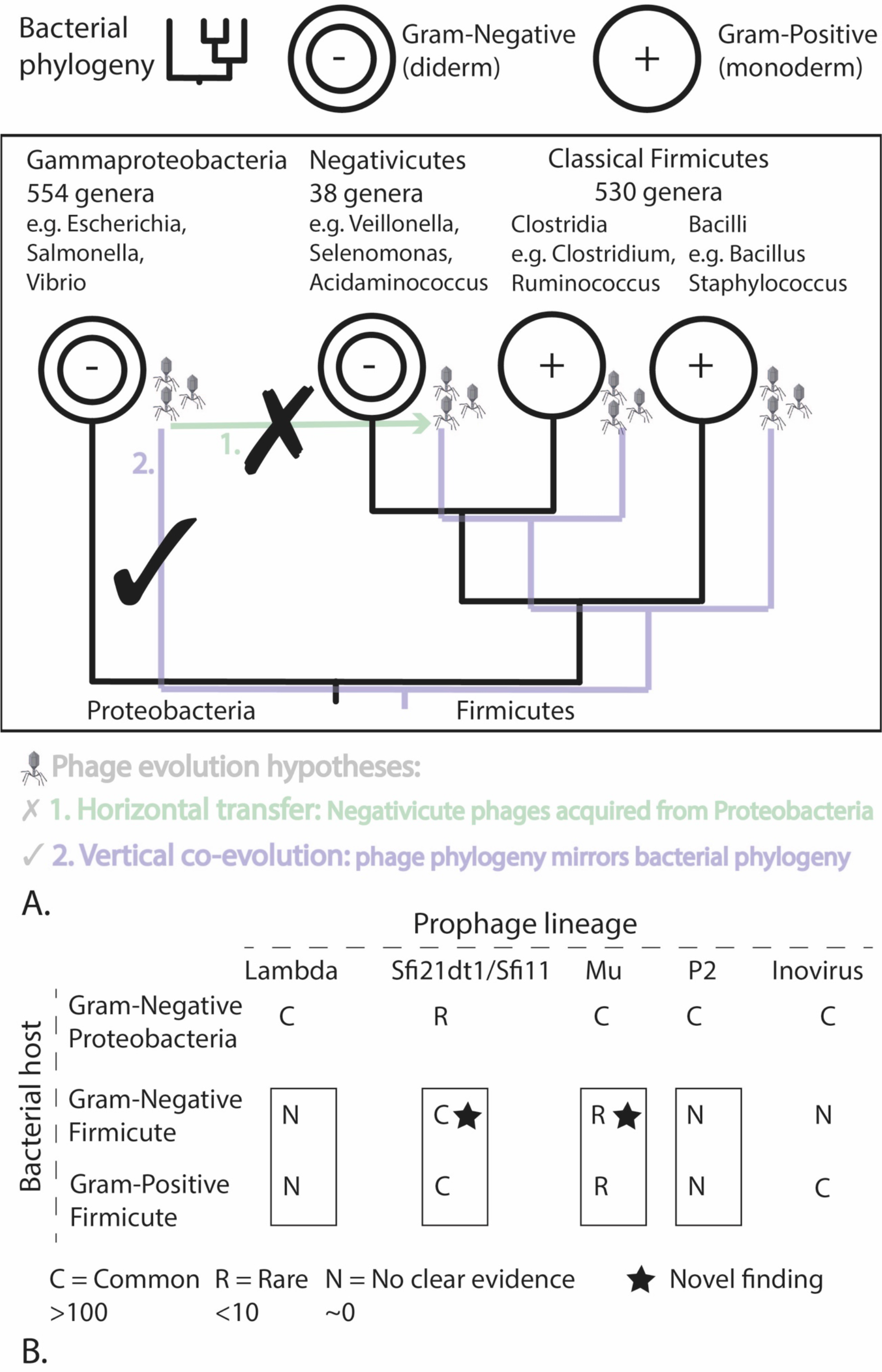
Phages of Gram-Negative Firmicutes are grouped with phages of Gram-Positive Firmicutes. **A**. Depiction of two contrasting hypotheses concerning the evolution of Negativicute phages. Our results support the phage-host co-evolution hypothesis. **B.** Summary of the different types of prophages putatively detected in different bacterial lineages.

## Materials and methods

### Annotation of prophage sequences

307 Negativicute and 43 Halanaerobiales genome sequences were downloaded from NCBI GenBank July 2018 (Clark et al., 2016). A further 200 complete genomes from each of the *Bacillus* and *Escherichia*, and 136 complete genomes for *Clostridium*, were also obtained from GenBank. The sequences were input to PHASTER (Arndt et al., 2016) via the API to predict candidate prophage sequences with the metagenomic contigs option selected to handle incomplete genome assemblies. The processing of data was performed in Python with the Biopython (Cock et al., 2009) and Snakemake modules (Koster and Rahmann, 2018) calling the following external tools. Open-reading frames were predicted in these possible prophage sequences with prodigal v2.6 (Hyatt et al., 2010). These protein sequences where then annotated with pairwise alignment against the large NCBI NR protein database with diamond v0.8.22 (Buchfink et al., 2015) and separately via profile HMM searches with HMMER3 v3.1b2 (HMMER, 2017) against the Prokaryotic Viral Orthologous Groups (pVOGS) (Grazziotin et al., 2017). We computed all alignment hits with e-value thresholds 0.1 and then for each protein we produced an initial consensus annotation by taking the top NR and pVOG hits, excluding annotations that were from hypothetical, unknown, or uncharacterised proteins.

We used the protein annotations to define the phage genome modules as follows with the corresponding associated protein annotations listed in parentheses: head (head, capsid, Clp protease, prohead protease), neck (head tail), portal (portal), tail (tail, tail fibre, tail tube, tail sheath, tail tape measure, base plate), packaging (terminase), replication (helicase, primase), lysis (lysis, lysin, holin), lysogeny (integrase, repressor, antirepressor). tRNAs were also annotated separately. For tail proteins, a more specific annotation was preferred over “tail” in the case of multiple hits. Any prophage sequences that contained fewer than two annotated proteins were discarded at this stage. Several *Escherichia* and *Bacillus* phages were also discarded after manual inspection as it was unclear if they represented individual phages or two adjacent. For the two *A. intestini* phages that we identified previously (Rands et al., 2018), we manually curated the prophage boundaries.

Prophage genome maps were visualized via R scripts using the ggplot2 and Gviz packages (Hahne and Ivanek, 2016). We aligned and visually inspected the genome maps to identify lambdoid phages and we searched for Mu and P2 phages systematically by looking for prophages with at least two annotated proteins relating to Mu-like or P2-like phages and then performed further manual inspection of the resulting prophage genome maps.

### Annotation of bacterial genes within prophages

Bacterial genes were tentatively annotated using the NR annotations as 16S, 23S, 30S, and 50S related proteins. Candidate antibiotic resistance genes were also preliminarily annotated based on NR protein hits relating to the following antibiotic categories: beta-lactam, aminoglycoside, chloramphenicol, glycopeptide, quinolone, tetracycline, macrolide, ansamycin, strepogramin, lipopeptide, vanomycin and efflux pump. Then all Gram-Negative Firmicute prophage region proteins were scanned using diamond with >=80% sequence identity and query coverage thresholds against a merged non-redundant database created in February 2018 of CARD protein homolog models (Jia et al., 2017) and sequences from Resfinder and ResfinderFG (Zankari et al., 2012). A single discrepancy was found (see Results and Discussion): three Metallo Beta-lactamases (MBLs) were annotated based on high hits (>=90% amino acid identity and query coverage) against an MBL fold metallo-hydrolase annotated protein in NR (accession: WP_084644750.1), but this MBL was not present in (and did not have a close homolog in) the merged database of curated ARG sequences.

Similarly, possible bacterial virulence factors, including toxins, were initially tentatively annotated based on NR, but Gram-Negative Firmicute phage sequences were also scanned for virulence factors by aligning the proteins against the VirulenceFinder database (Joensen et al., 2014) with diamond v0.8.22 and 80% sequence identify and query coverage thresholds. NR virulence factor hits that were not confirmed by the VirulenceFinder search were merely annotated as likely bacterial genes, with a few exceptions: Virulence associated E-proteins (VapE) showed homology to phage primases and helicases and the phage replication annotation was preferred in these cases; bacterial toxins sometimes show homology to phage holins, thus annotated toxins co-annotated as holins were resolved to phage holins. Transposon related genes were annotated based on transposases and recombinases in NR.

### Clustering of prophage genome sequences

The 3,462 full-length filtered prophage genome sequences and the annotated portal protein sequences were clustered with CD-HIT and PSI-CD-HIT v4.7 (Fu et al., 2012) at 80% and 40% sequence identify cut-offs. The subsequent clusters were visualized via the Python matplotlib and networkx modules. The sequences within the cluster containing both Veillonallales and Acidaminococcales phages were aligned together to create a multiple sequence alignment with MAFFT v7.309 (Katoh and Standley, 2013) with 1000 iterations and the adjustdirectionaccurately flag. We then built Maximum Likelihood phylogenies with iqtree v1.1.5 (Nguyen et al., 2015) using 1000 UFBoot2 ultrafast bootstraps (Hoang et al., 2018) and the HKY DNA model. We visualised the phylogenetic trees with Jalview (Waterhouse et al., 2009).

### Identification of extracellular phages in human gut viromes

183 human gut viromes (sequences of viral-like particles; VLPs) were downloaded from BioProject PRJEB7772 (171 samples) (Norman et al., 2015) and PRJNA356544 (12 samples) (Minot et al., 2012). Paired end Illumina derived sequencing reads from both studies were pre-processed to remove low quality and artefactual sequences with trimmomatic v0.38 and parameters -phred33 ILLUMINACLIP:TruSeq3-PE.fa:2:30:10 LEADING:3 TRAILING:3 SLIDINGWINDOW:4:15 MINLEN:36 (Bolger et al., 2014). Next, low complexity sequences were removed with tagDust2 v2.33 (Lassmann, 2015) with a dust threshold of 16.

Reads were aligned to the reference bacterial genomes with bowtie v2.3.2 (Langmead and Salzberg, 2012), and subsequently BAM files were processed with SAMtools v1.8 (Li et al., 2009) to remove low quality mapping with MAPQ 10 threshold. Next bedtools v2.25 (Quinlan and Hall, 2010) and matplotlib were used to generate the coverage plots, pooling the results across all samples.

To identify the *V. parvula* extracellular phages, the reads were assembled with MEGAHIT v1.1.3 (Li et al., 2015) with k-min 27. Upon assembly it became clear that the predicted *V. parvula* prophage 1 was truncated and so the prediction was extended to include the full prophage sequence. Sequences were rotated with MARS (Ayad and Pissis, 2017) to ensure that the sequences started and ended at the same point (since the genomes are circular), and global alignment was performed with emboss stretcher (Rice et al., 2000) with gapopen 16 and gapextend 4 penalties. Percent matches between the sequences was then calculated and genome maps visualized.

## Acknowledgements

We thank Konstantin Popadin, Elizaveta Starikova, Francisco Brito, and Jennifer Tan for helpful comments. We are funded by SNSF grant IZLRZ3_163863. We are grateful to the Baobab HPC cluster at the University of Geneva for computational resources. The authors declare no conflicts of interest.

