## Supplementary material for "Comparative genomics groups phages of Negativicutes and classical Firmicutes despite different Gram-staining properties"

### Supplementary Figure legends for Rands *et al.* 2019

**Figure S1:** Display of all 485 prophage genome maps predicted from Gram-Negative Firmicutes. Each horizontal line corresponds to an individual prophage shown to scale and color-coded for annotated phage genes according to the key displayed in the right-side box. The left vertical bar indicates the bacterial host in a colour code.

**Figure S2:** Projection of virome sequences from 183 human stool samples on **A.** *Acidaminococcus intestini* RYC-MR95, and **B.** *Veillonella parvula* UTDB1-3. The first panel shows the read coverage (Y-axis) across the complete bacterial genome sequence (X-axis; with bp coordinates). Predicted prophage regions are marked with red triangles and magnified in the subsequent panels. Virome reads projected outside of prophage prediction are listed in **Table S4**.

**Figure S3**: The same display of virome sequences projected onto bacterial genomes as in **Figure S2**, but for two different Negativicute species: **A.** *Dialister Marseille*, and **B.** *Negativicoccus massiliensis*. For non-phage peak annotations, see **Table S4**.

**Figure S4:** Gene flanking analysis for the lysis module from all prophages predicted in all the different bacterial clades (**Table S2**), a total of 3,462 prophages. The lysis module is generally located next to the tail module in Firmicute prophages, but adjacent to the packaging (terminase) module in Escherichia phages.

**Figure S5**: Candidate Mu-like prophage in the Negativicute *Propionispora vibrioides*. Phage-related genes (arrows indicate transcription direction) are coloured and show characteristics of Mu-like genome organization.

**Figure S6**: The genome maps of Negativicute prophages harbouring candidate antibiotic resistance genes MBL (top three Veillonella prophages) and tet(32) (bottom Selenomonas prophage remnant); excludes the ACI-1 prophage harbouring example characterised previously (Rands et al., 2018). The MBL gene was annotated based on a high match (>90% sequence identity and query coverage) to an MBL fold metallo-hydrolase annotated protein with protein accession WP_084644750.1. Note that this gene also has non-ARG functions. The tet(32) gene was also annotated in NR with protein accession WP_002602099.1 (with 100% sequence identity and 99% query coverage), and was also present in the ResfinderFG database with the annotation tetracycline protection.

**Figure S7**: Prophage gene annotation versus the prophage genome size. The X-axis shows the prophage genome size in bp and the Y-axis is the percentage of the given annotation specified in the colour code for each prophage (discarding proteins without annotation within these modules). If prophages show directionality in the process of gene deletion when becoming prophage remnants, then we expect an increased prevalence of the preserved gene types and hence a higher percentage of these at smaller prophage genome size. The equations and $R^2$ values of the linear regression lines are shown and give an indication that the data is variable without a clear negative or positive correlation suggesting random deletion of prophage modules. Data shown for **A.** all prophages and **B.** prophages from complete bacterial genome assemblies.

### Supplementary Figures

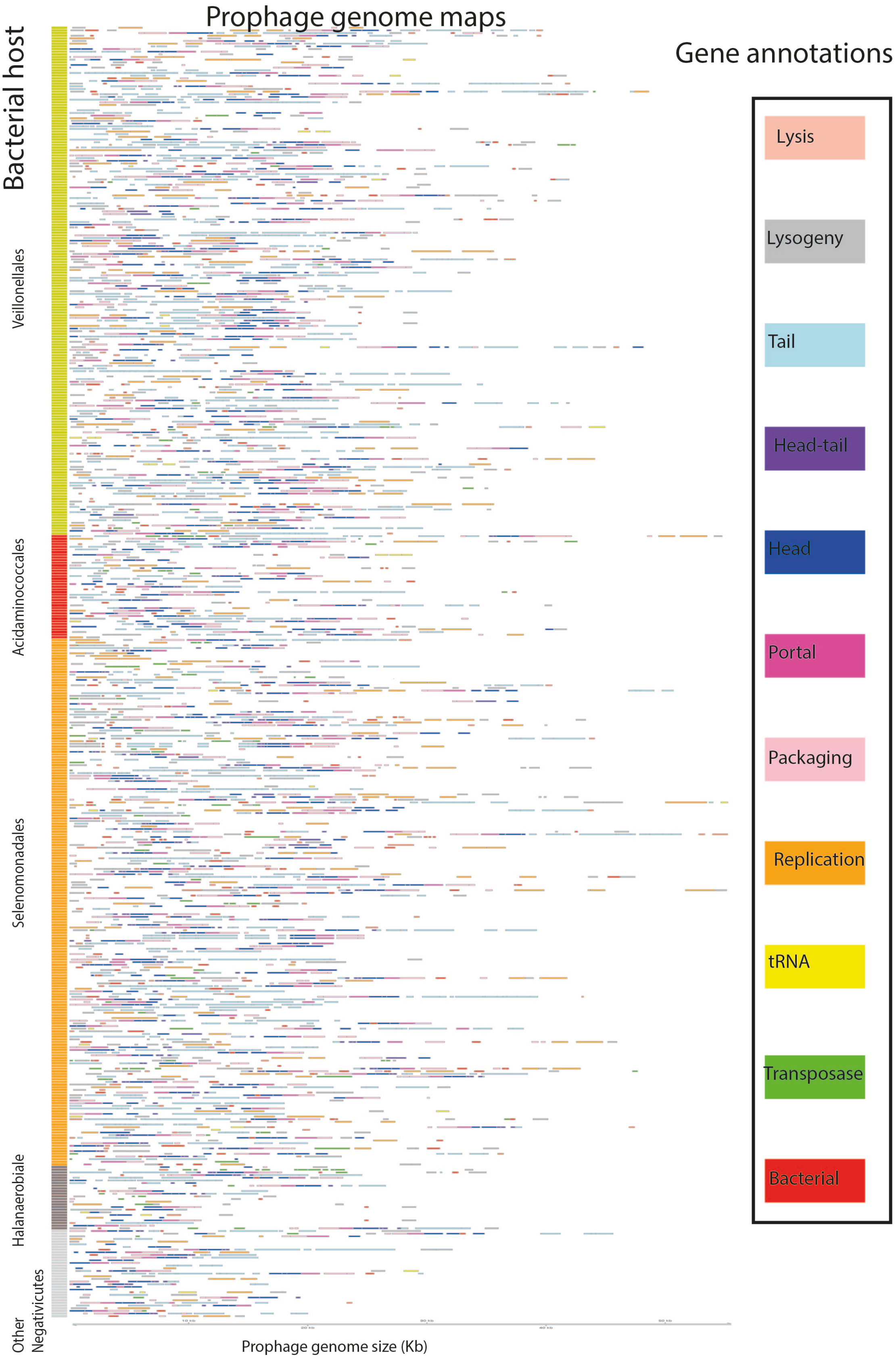

**Figure S1**

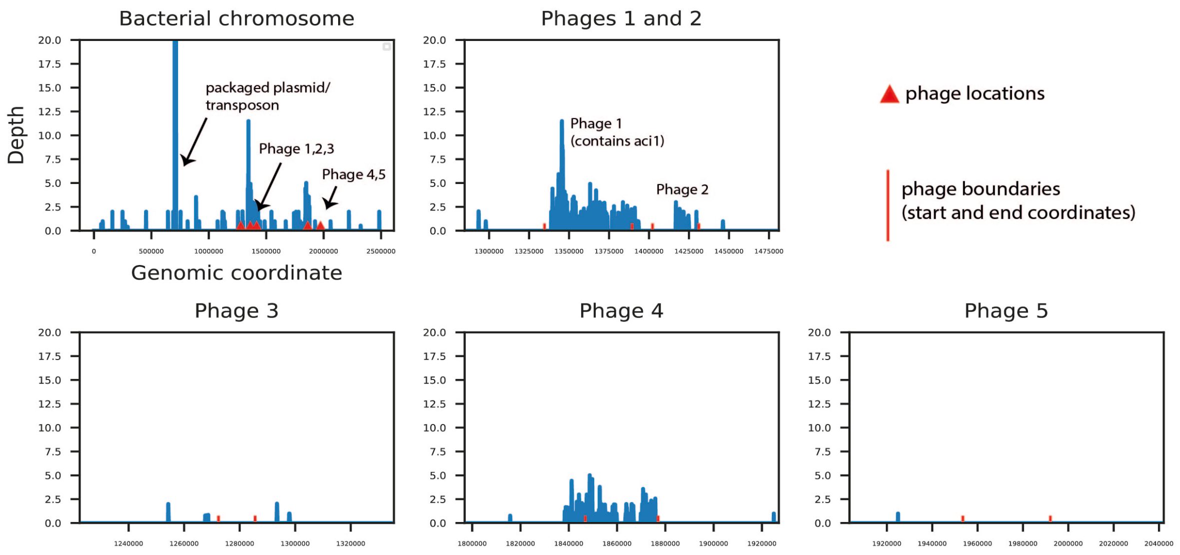

A. Acidaminococcus intestini RYC-MR95

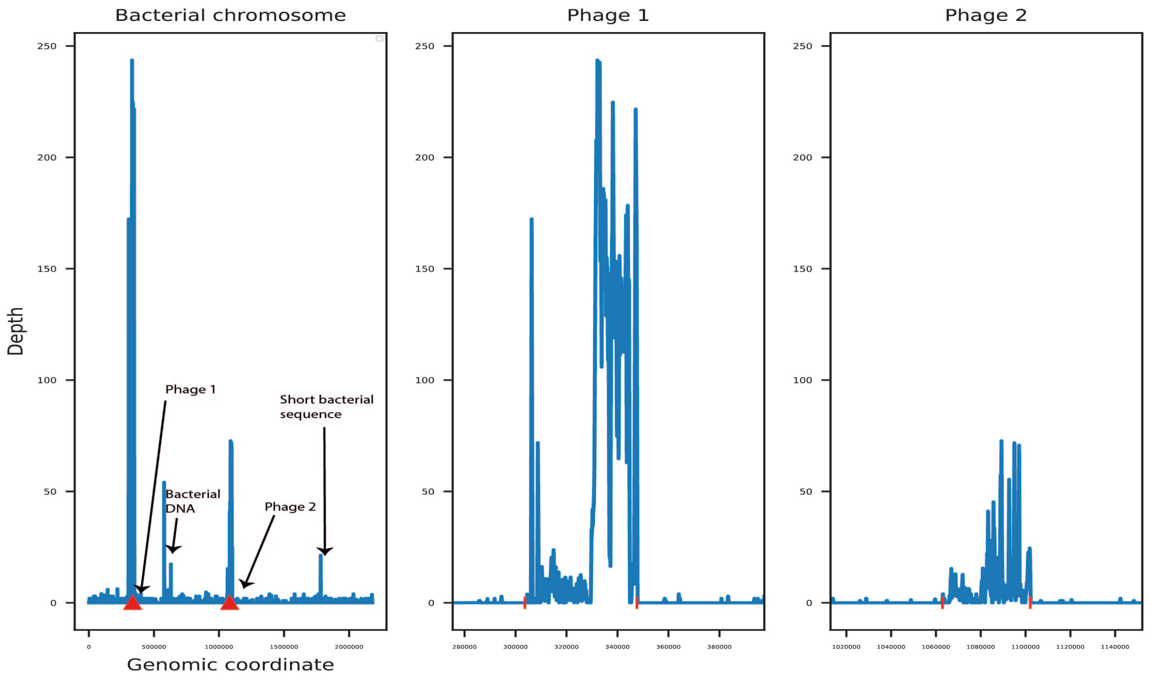

B. Veillonella parvula UTDB1-3

**Figure S2**

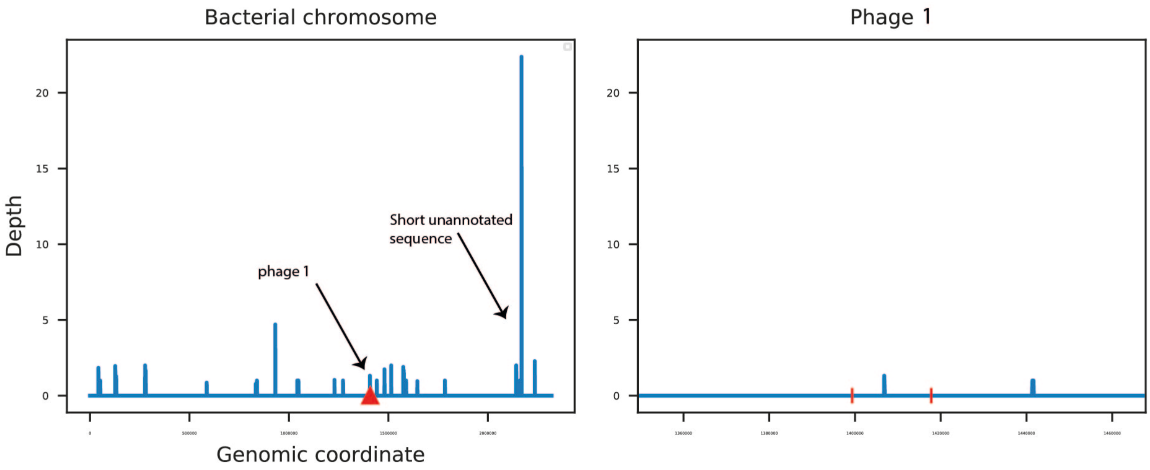

A. Dialister sp. Marseille-P5638

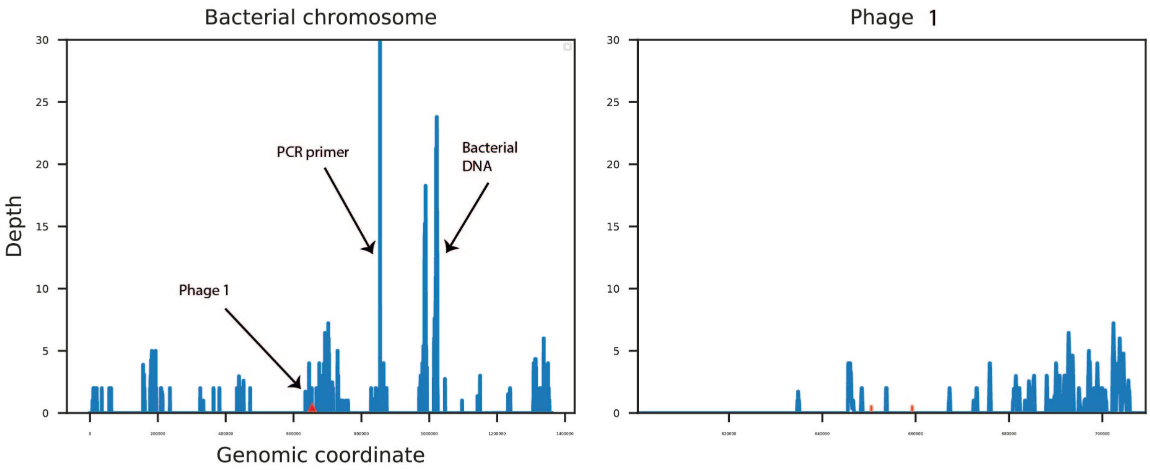

B. Negativicoccus massiliensis Marseille-P2082

phage locations

phage boundaries (start and end coordinates)

**Figure S3**

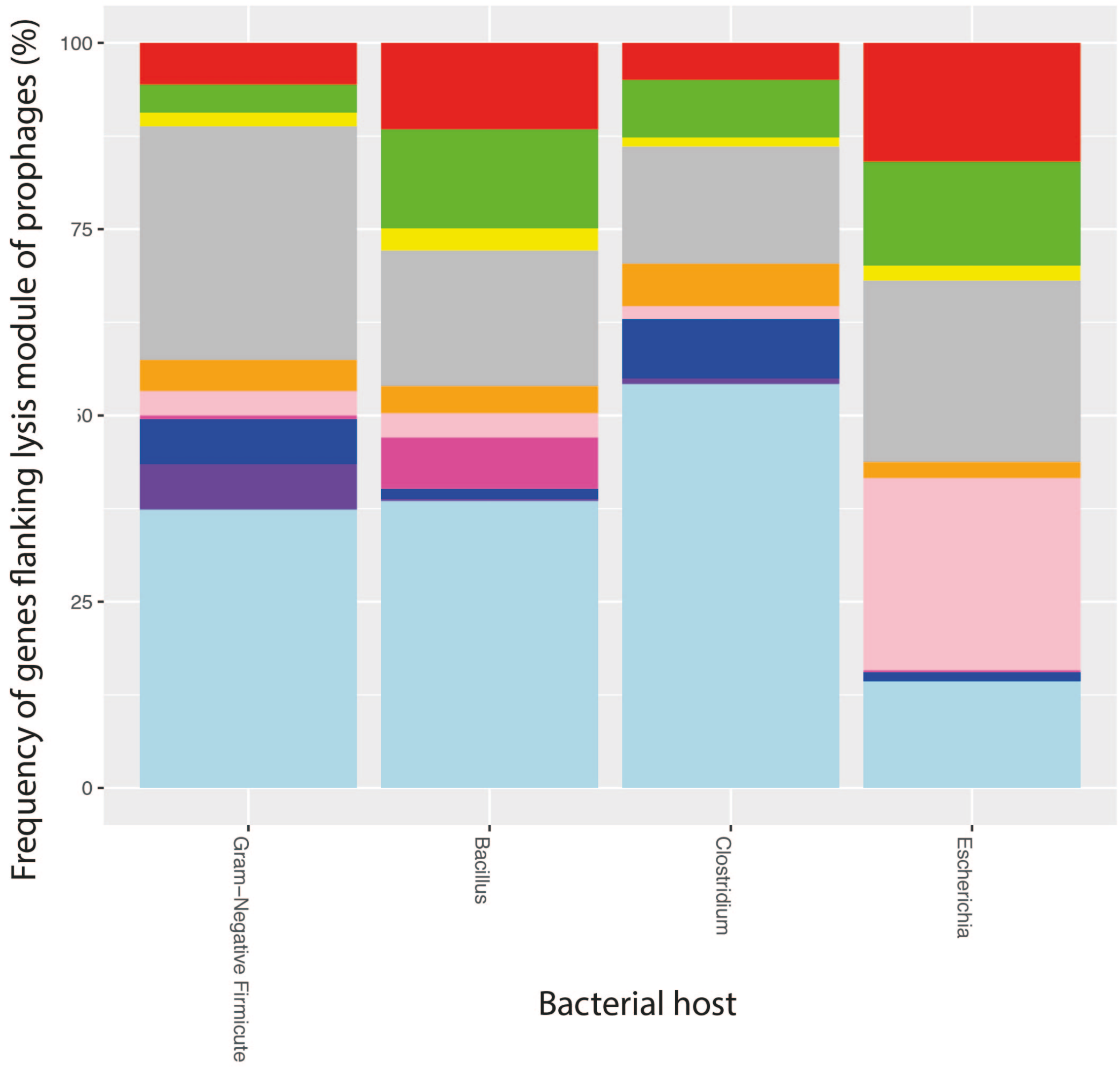
Frequency of genes flanking lysis module of prophages (%)
100
75
50
25
0
Gram−Negative Firmicute
Bacillus
Clostridium
Escherichia
Bacterial host

**Figure S4**

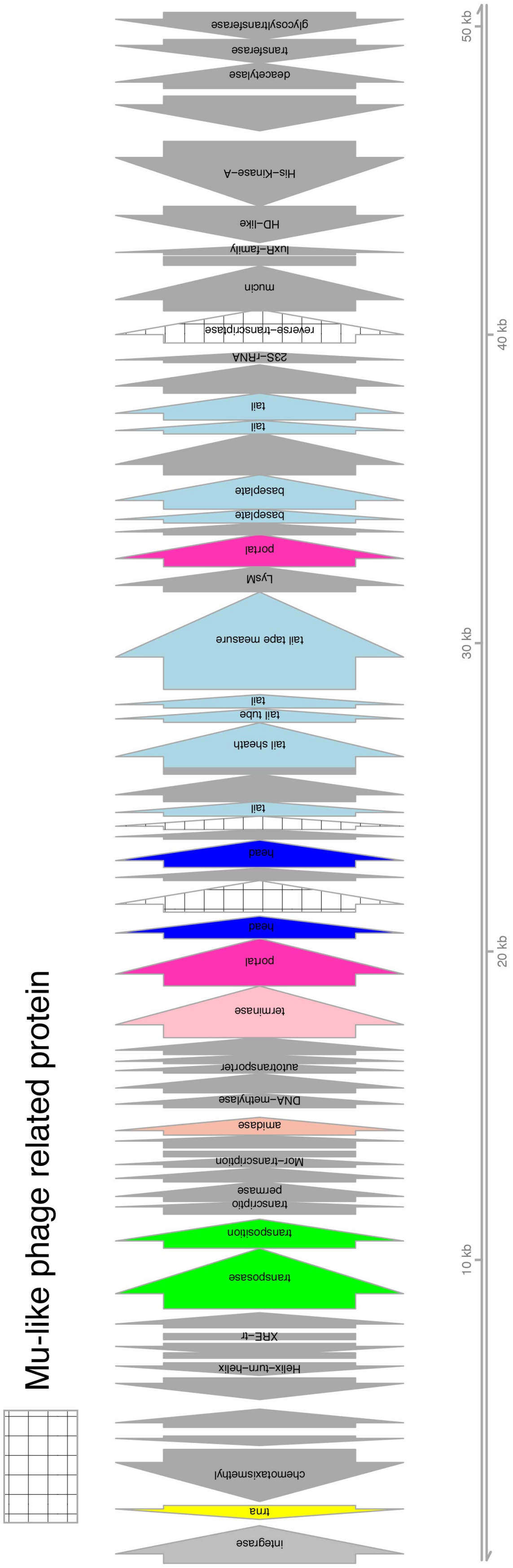
Mu-like phage related protein
integrase
trna
chemotaxismethyl
Helix−turn−helix
XRE−tr
transposase
transposition
transcriptio
permase
Mor−transcription
amidase
DNA−methylase
autotansporter
terminase
portal
head
head
tail
tail sheath
tail tube
tail
tail tape measure
LysM
portal
baseplate
baseplate
tail
tail
23S−rRNA
reverse−transcriptase
mucin
luxR−family
HD−like
His−Kinase−A
deacetylase
transferase
glycosyltransferase
10 kb
20 kb
30 kb
40 kb
50 kb

**Figure S5**

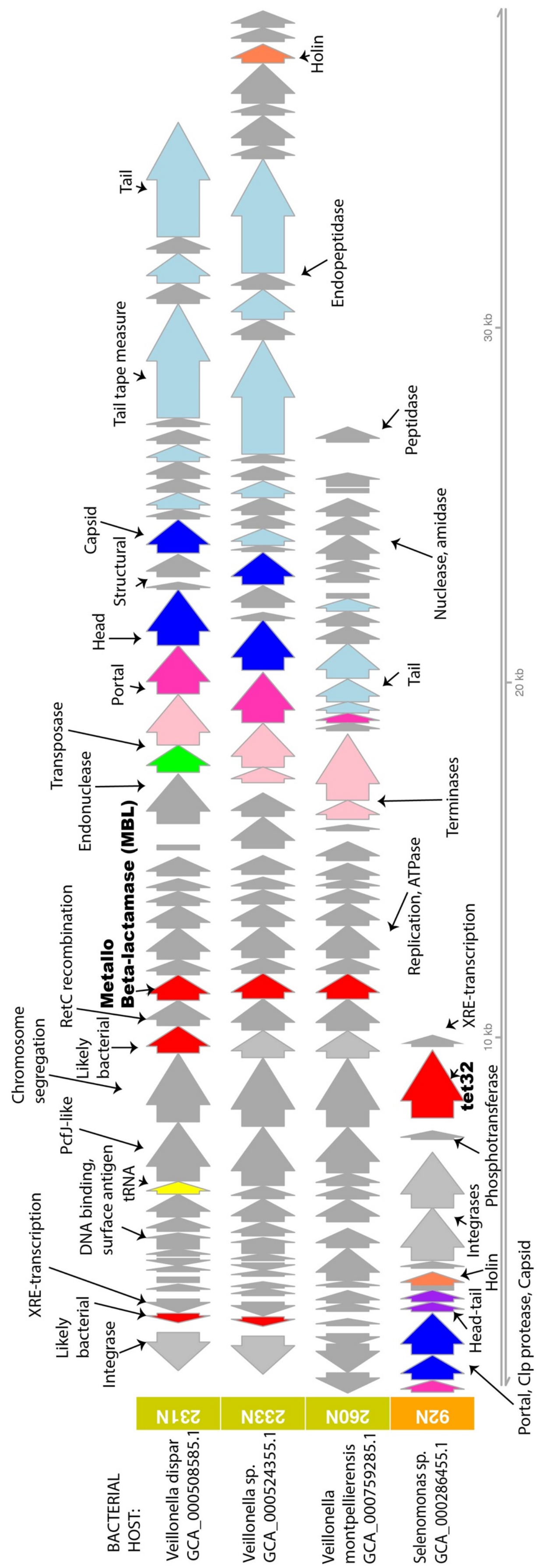
BACTERIAL HOST:
Veillonella dispar GCA_000508585.1
Veillonella sp. GCA_000524355.1
Veillonella montpellierensis GCA_000759285.1
Selenomonas sp. GCA_000286455.1
231N
233N
260N
92N
Integrase
Likely bacterial
XRE-transcription
DNA binding, surface antigen
tRNA
PcfJ-like
Chromosome segregation
Likely bacterial
RetC recombination
Metallo Beta-lactamase (MBL)
Endonuclease
Transposase
Portal
Head
Structural
Capsid
Tail tape measure
Tail
Endopeptidase
Holin
Replication, ATPase
Terminases
Tail
Nuclease, amidase
Peptidase
Portal, Clp protease, Capsid
Head-tail
Holin
Integrases
Phosphotransferase
tet32
XRE-transcription
10 kb
20 kb
30 kb

**Figure S6**

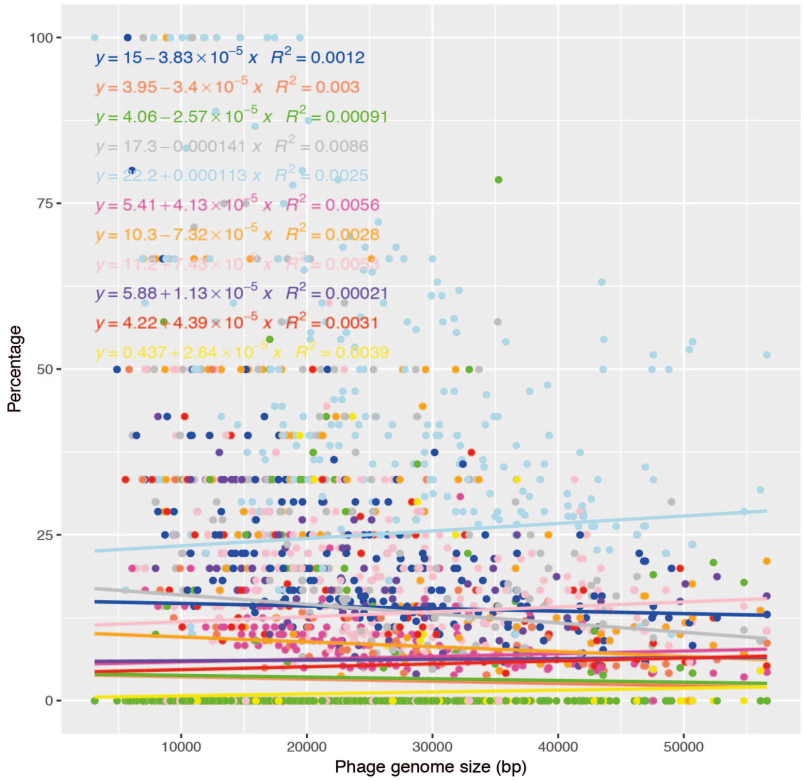

A. Prophages in all bacterial genomes

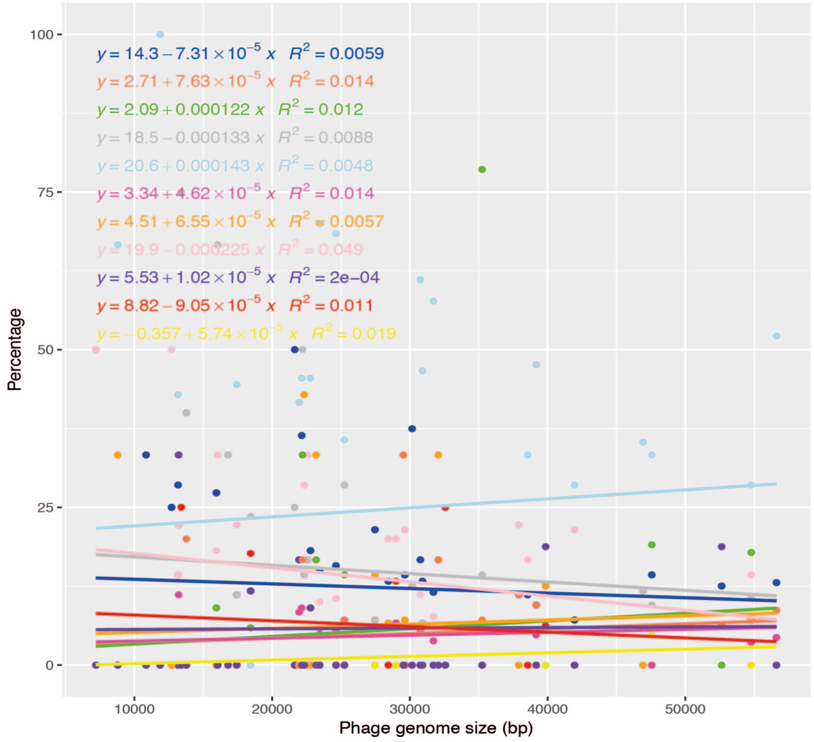

B. Prophages in complete bacterial genomes

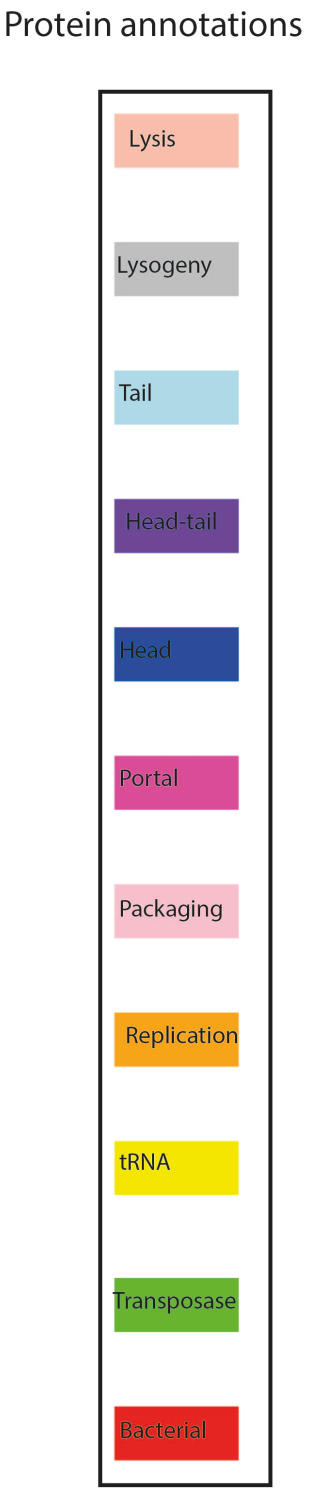

**Figure S7**

### Supplementary Tables

| **Bacterial host** | **Bacterial assemblies** | **Prophages** | **Number of prophages per bacterial genome** |
|---|---|---|---|
| *Megasphaera* | 50 | 115 | 2.3 |
| *Veillonella* | 47 | 44 | 0.9 |
| *Selenomonas* | 42 | 80 | 1.9 |
| Other Negativicute | 30 | 31 | 1 |
| *Halanaerobium* | 27 | 9 | 0.3 |
| *Acidaminococcus* | 20 | 26 | 1.3 |
| *Dialister* | 17 | 18 | 1.1 |
| *Megamonas* | 13 | 11 | 0.8 |
| *Sporomusa* | 9 | 37 | 4.1 |
| *Pelosinus* | 9 | 14 | 1.6 |
| *Phascolarctobacterium* | 8 | 11 | 1.4 |
| *Veillonellaceae* | 6 | 3 | 0.5 |
| *Selenomonadaceae* | 6 | 7 | 1.2 |
| *Mitsuokella* | 6 | 5 | 0.8 |
| *Anaerovibrio* | 6 | 11 | 1.8 |
| *Acidaminococcaceae* | 6 | 1 | 0.2 |
| *Sporomusaceae* | 5 | 6 | 1.2 |
| *Orenia* | 4 | 8 | 2 |
| *Candidatus* | 4 | 2 | 0.5 |
| *Selenomonadales* | 3 | 3 | 1 |
| *Halothermothrix* | 3 | 1 | 0.3 |
| *Succiniclasticum* | 2 | 1 | 0.5 |
| *Propionispora* | 2 | 4 | 2 |
| *Propionispira* | 2 | 4 | 2 |
| *Negativicoccus* | 2 | 1 | 0.5 |
| *Thermosinus* | 1 | 1 | 1 |
| *Succinispira* | 1 | 0 | 0 |
| *Sporolituus* | 1 | 0 | 0 |
| *Selenihalanaerobacter* | 1 | 0 | 0 |
| *Schwartzia* | 1 | 2 | 2 |
| *Negativicutes* | 1 | 1 | 1 |
| *Massilibacillus* | 1 | 5 | 5 |
| *Halonatronum* | 1 | 0 | 0 |
| *Halobacteroides* | 1 | 0 | 0 |
| *Halarsenatibacter* | 1 | 3 | 3 |
| *Dendrosporobacter* | 1 | 3 | 3 |
| *Colibacter* | 1 | 1 | 1 |
| *Centipeda* | 1 | 3 | 3 |
| *Caecibacter* | 1 | 3 | 3 |
| *Anaerosporomusa* | 1 | 0 | 0 |
| *Anaeromusa* | 1 | 6 | 6 |
| *Anaeroglobus* | 1 | 1 | 1 |
| *Anaeroarcus* | 1 | 1 | 1 |
| *Allisonella* | 1 | 1 | 1 |
| *Acetonema* | 1 | 0 | 0 |
| *Acetohalobium* | 1 | 1 | 1 |

**Table S1**: The distribution of prophages across different Gram-Negative Firmicute taxa.

| Clade | Number of complete bacterial genomes analysed | Number of prophages | Mean prophages per bacterial genome |
|---|---|---|---|
| Clostridium | 136 | 434 | 3.2 |
| Bacillus | 200 | 793 | 3.9 |
| Escherichia | 200 | 1750 | 8.8 |

**Table S2**: The number of prophages predicted and annotated in this study from bacterial genomes other than the Gram-Negative Firmicutes.

| Species/strain name | NCBI Identifiers | Prophages | Ecosystem | Reference or Accession |
|---|---|---|---|---|
| Halothermothrix oreni* | GCA_000020485.1., CP001098.1 | 1 | Salt lakes | (Cayol et al., 1994) |
| Acidaminococcus fermentans | GCA_000025305.1., CP001859.1 | 1 | Pig gut | (Chang et al., 2010) |
| Acetohalobium arabaticum* | GCA_000144695.1., CP002105.1 | 1 | Salt lakes | (Sikorski et al., 2010) |
| Halanaerobium hydrogeniformans* | GCA_000166415.1., .CP002304.1 | 1 | Alkali lakes | (Brown et al., 2011) |
| Selenomonas sputigena | GCA_000208405.1., CP002637.1 | 2 | Human mouth | (Nagpal et al., 2016) |
| Acidaminococcus intestini | GCA_000230275.1, .CP003058.1 | 5 | Human gut | (D'Auria et al., 2011) |
| Pelosinus fermentans | GCA_000271665.2., .CP010978.1 | 2 | Ground water | (Bowen De Leon et al., 2012) |
| Selenomonas ruminantium | GCA_000284095.1., AP012292.1, AP012300.1, AP012301.1 (chromosome, 2 plasmids) | 4,<br>1,<br>1 | Sheep gut | (Kaneko et al., 2015) |
| Pelosinus sp. UFO1 | GCA_000725345.1., CP008852.1 | 1 | Soil | (Brown et al., 2014) |
| Megasphaera elsdenii 14-14<br>NCIMB702410<br>DSM_20460 | GCA_001304715.1., CP009240.1,<br>GCA_003006415.1/ CP027569.1,<br>GCA_003010495.1/ CP027570.1 | 2,<br>4,<br>1 | Cow/sheep guts | (Marx et al., 2011) |
| Selenomonas sp. oral taxon 136,<br>920,<br>478 | GCA_001554015.1/ CP014239.1,<br>GCA_001717585.1/ CP017042.1, CP017043.1 (chr, plasmid)<br>GCA_001189555.1,CP012071.1 | 3,<br>4,<br>1,<br>0 | Human mouth | (Dewhirst et al., 2010) |
| Veillonella parvula UTDB1-3<br>DSM 2008<br>NCTC11810 | GCA_002005185.1, CP019721.1,<br>GCA_000024945.1, CP001820.1<br>GCA_900186885.1,LT906445.1 | 2,<br>0,<br>0 | Human gut/ mouth/ vagina | (Bhatti and Frank, 2000); and others |
| Megasphaera hexanoica | GCA_003315775.1, CP011940.1 | 7 | Cow gut | (Jeon et al., 2017) |
| Veillonella rodentium | GCA_900187285.1, LT906470.1 | 1 | Hamster mouth | Biosample SAMEA44547418 |
| Dialister sp. Marseille | GCA_900343095.1, LT996885.1 | 1, | Human gut | Bioproject PRJEB25867 |
| Dialister pneumosintes strain F0677 | GCA_001717505.1,CP017037.1 | 0 | Human nose/mouth | (Rousee et al., 2002) |
| Veillonella atypica strain OK5 | GCA_002082765.1,CP020566.1 | 0 | Human mouth | (Egland et al., 2004) |
| Megasphaera sp. AJH120 | GCA_003367905.1,CP029462.1 | 0 | Chicken gut | (Maki and Looft, 2018) |
| Negativicoccus massiliensis strain AT7 | GCA_900155405.1,LT700188.1 | 1 | Human gut | (Togo et al., 2016) |
| Megamonas hypermegale strain NCTC10570 | GCA_900187035.1,LT906446.1 | 0 | Chicken gut | (Morotomi et al., 2007) |

**Table S3**: The complete bacterial genomes of Gram-Negative Firmicutes (Negativicutes and Halanaerobiales*) with the number of predicted prophages and the ecosystem they inhabit.

| ORF NUMBER | ANNOTATION | DATABASE SOURCE |
|---|---|---|
| **Peak: 702475- 709975bp in A. intestini** | | |
| 1 | Tet32/TetM/TetW/TetO/TetS | NR |
| 2 | XRE family transcriptional regulator | NR, pVOGs |
| 3 | methyltransferase | NR, pVOGs |
| 4 | sigma-70 family RNA polymerase sigma factor | NR, pVOGs |
| 5 | helix-turn-helix domain-containing protein | NR |
| 6 | plasmid mobilization relaxosome protein MobC | NR |
| 7 | relaxase | NR |
| **Peak: 577675-630475bp in V. parvula** | | |
| 3 | DNA/RNA nuclease SfsA | NR |
| 7 | Cro/CI family transcriptional regulator | NR |
| 8 | Nitroreductase family protein | NR |
| 9 | OmpH family outer membrane protein | NR |
| 10 | sporulation protein YtfJ | NR |
| 11 | CsrA-like transcriptional regulator | NR |
| 13 | adenosyltransferase | NR |
| 14 | M23 family metallopeptidase | NR |
| 15 | terminal repeat-encoded protein | pVOGs |
| 16 | Putative GTP-binding protein | NR |
| 17 | Chromosome (plasmid) partitioning ParB | NR |
| 18 | ParA family | NR |
| 19 | methionine synthase | NR |
| 20 | homocysteine methyltransferase | NR |
| 21 | methylenetetrahydrofolate reductase | NR |
| 22 | rRNA small subunit methyltransferase | NR |
| 23 | chaperone protein DnaK | pVOGs |
| 24 | tRNA synthesis enzyme | NR |
| 25 | GTPase / small terminase | NR/ PVOGs |
| 27 | membrane protein insertase YidC | NR |
| 28 | ribonuclease P | NR |
| 29 | 50S ribosomal protein L34 | NR |
| 30 | chromosomal replication initiator dnaA | NR |
| 31 | DNA polymerase III subunit beta | NR |
| 32 | RNA-binding S4 domain-containing | NR |
| 33 | DNA replication/repair RecF | NR |
| 36 | topoisomerase subunit B | NR |
| 37 | amino acid permease | NR |
| 38 | YbjQ family | NR |
| 39 | efflux RND transporter permease | NR |
| 40 | Putative RND family drug transporter | NR |
| 41 | HlyC/CorC family transporter | NR |
| 42 | SAM-dependent methyltransferase | NR |
| 43 | Transcriptional regulator ATRX-like | NR |
| **Peak: 1781325-1781825bp in V. parvulla** | | |
| 1 | RNA polymerase sporulation sigma factor SigH | NR, pVOGs |
| **Peak: 2168675-2168925bp in D. Marseille (1 unannotated protein/nucleotide sequence only)** | | |
| **Peak: 854675-854725bp in N. massiliensis** | | |
| 1 nucleotide sequence | Illumina PCR Primer | VecScreen |

| Peak: 986125-1024125 in N. massiliensis | | |
|---|---|---|
| 1 | aspartate 1-decarboxylase | NR |
| 2 | pantoate beta-alanine ligase | NR |
| 3 | bile acid:sodium symporter family | NR |
| 5 | MFS transporter | NR |
| 6 | DNA gyrase subunit A | NR |
| 7 | RepB family plasmid replication initiator | NR |
| 9 | Rz1 | pVOGs |
| 10 | magnesium transporter | NR |
| 11 | 30S ribosomal protein S6 / internal head protein | NR / pVOGs |
| 12 | single-stranded DNA-binding protein | NR, pVOGs |
| 13 | 30S ribosomal protein S18 | NR |
| 15 | YadA-like C-terminal domain | NR |
| 16 | TetR/AcrR family transcriptional regulator | NR |
| 17 | cation-translocating P-type ATPase | NR |
| 19 | nickel-dependent lactate racemase | NR |
| 20 | pyridoxamine 5'-phosphate oxidase | NR |
| 21 | Hydrolase of the alpha/beta family | NR |
| 22 | ribonucleoside-triphosphate reductase | NR, pVOGs |
| 23 | haloacid dehalogenase-like hydrolase | NR |
| 26 | band 7-like protein | pVOGs |
| 27 | nodulation protein NfeD/ Clp protease | NR, pVOGs |
| 28 | Somatostatin | NR |
| 30 | cobaltochelatase subunit CobN | NR |
| 31 | mechanosensitive ion channel | NR |
| 32 | ABC transporter permease subunit | NR |
| 33 | ABC transporter permease subunit | NR |
| 34 | ABC transporter ATP-binding protein | NR |
| 35 | energy-coupled thiamine transporter ThiT | NR |

**Table S4:** Annotations for regions of non-prophage regions with peaks of coverage based on read mapping from VLPs derived from the human gut samples; see the corresponding **Figures S2 and S3**. Proteins without annotations are omitted.
